# Lysosome transporter purification and reconstitution identifies Ypq1 pH-gated lysine transport and regulation

**DOI:** 10.1101/2023.03.31.535002

**Authors:** Felichi Mae Arines, Aleksander Wielenga, Olive E. Burata, Francisco Narro Garcia, Randy B. Stockbridge, Ming Li

## Abstract

Lysosomes achieve their function through numerous transporters that import or export nutrients across their membrane. However, technical challenges in membrane protein overexpression, purification, and reconstitution hinder our understanding of lysosome transporter function. Here, we developed a platform to overexpress and purify the putative lysine transporter Ypq1 using a constitutive overexpression system in protease- and ubiquitination-deficient yeast vacuoles. Using this method, we purified and reconstituted Ypq1 into proteoliposomes and showed lysine transport function, supporting its role as a basic amino acid transporter on the vacuole membrane. We also found that the absence of lysine destabilizes purified Ypq1 and causes it to aggregate, consistent with its propensity to be downregulated *in vivo* upon lysine starvation. Our approach may be useful for the biochemical characterization of many transporters and membrane proteins to understand organellar transport and regulation.

## Introduction

The lysosome is important in cellular homeostasis due to its role in the recycling of cellular material, the sensing of nutrient conditions, and the modulation of metabolic pools in response to nutrient excess or starvation. These are achieved, in part, through nutrient transporters on the lysosome membrane. After the degradation of cellular materials through endocytosis or autophagy, amino acid transporters export free amino acids to the cytosol to be re-used for protein synthesis. In yeast, the storage function of the vacuole (yeast lysosome) relies on importers to uptake and accumulate amino acids, which are later mobilized during starvation (Li and Kane, 2009; Lim and Zoncu, 2016). Metabolic sensing is also linked to lysosomal transporters, such as in arginine and cholesterol sensing through SLC38A9 and its connection to the mTORC1 pathway (Wang et al., 2015; Wyant et al., 2017), and in basic amino acid sensing through PQLC2 (Gasnier, 2021).

Lysosomal transporters, therefore, have an emerging prominence in our understanding of cellular regulation. Despite their importance, only a handful of lysosome membrane transporters have been biochemically characterized so far due to difficulties in membrane protein expression and purification (Lei et al., 2018; Tomita et al., 2021; Wang et al., 2015; Wyant et al., 2018; Zhang et al., 2020). First, membrane proteins are typically expressed at low levels, thus requiring overexpression and the use of large purification cultures. Eukaryotic membrane proteins rarely express well in heterologous bacterial systems, since doing so abolishes functionally important post-translational modifications. Second, membrane proteins often suffer instability upon removal from the lipid bilayer and solubilization with an imperfect membrane mimetic such as detergents. As a result, most lysosomal transporters (dozens in yeast and hundreds in humans) remain uncharacterized (Xiong and Zhu, 2016; Xu and Ren). We do not know their substrates, transport directionality, or energy dependence, among other properties.

There have been some efforts to understand lysosomal basic amino acid transport using genetic and other indirect methods. For example, transporter activity has been evaluated using purified vacuoles, in heterologous systems using oocytes, and in the presence of toxic amino acid analogs (Cools et al., 2020; Jezegou et al., 2012; Kawano-Kawada et al., 2019; Leray et al., 2021; Manabe et al., 2016; Sekito et al., 2014). These studies have provided great insights into transporter function but could be prone to controversy due to the indirect nature of these systems. Purified vacuoles and oocytes may not be a clean system for measuring transporter activity because of the functional redundancy of multiple transporters. In addition, transporters on the vacuole are open to regulatory cross-talk – deletion of one transporter gene can sometimes affect the transcription and stability of other transporter genes (MacDiarmid et al.; Velivela and Kane, 2018). Thus, heterologously expressed or deletion-based *in vivo* and semi-*in vitro* transport assays could be hard to interpret.

Thus, there is an urgent need for accessible methods to express, purify, and reconstitute lysosomal transporters into liposomes for biochemical characterization. Here, we developed a pipeline to express, purify, and reconstitute Ypq1 into proteoliposomes for functional studies. With this method, we showed lysine transport by Ypq1 in a reconstituted system. Increased uptake of radiolabelled lysine under a pH gradient implies that Ypq1 functions as a lysine/proton antiporter. Considering the low pH within the vacuole, our data do not support Ypq1 to be an exporter of positively charged amino acids (Jezegou et al., 2012); rather, our experiments imply that Ypq1 functions as an importer. Strikingly, purified Ypq1 depends on lysine (or arginine) for its stability *in vitro,* consistent with our *in vivo* observation that lysine withdrawal can trigger the ubiquitination and degradation of Ypq1 (Arines et al., 2021; Li et al., 2015b; Zhang and Ye, 2021). Furthermore, this lysine/arginine dependence is abolished in a degradation-resistant PQ-motif mutant, and exacerbated in a constitutively-degrading M73D mutant, suggesting a direct link between protein stability and substrate-dependent regulation. Our method is adaptable to other eukaryotic transporters and has the potential to support the field in systematically characterizing transporters on the lysosome and other organelles.

## RESULTS

### Constitutive overexpression in rich media maximized Ypq1 expression

To overexpress Ypq1 for purification and functional studies, we tested the effect of galactose-inducible and constitutive overexpression systems on Ypq1-GFP protein levels (Fig. 1A-B). For galactose induction, we cloned YPQ1 CDS into the pDDGFP2-LEU2d vector and transformed the construct into BJ2168 yeast lacking the major vacuolar protease Pep4 (Drew et al., 2008; Parker and Newstead, 2014). For constitutive overexpression, we cloned YPQ1-GFP CDS into integration plasmids containing the ADH1, GPD1, or TEF1 promoters and stably integrated these plasmids into the genome of SEY6210 *PEP4Δ* yeast.

**Fig. 1.**
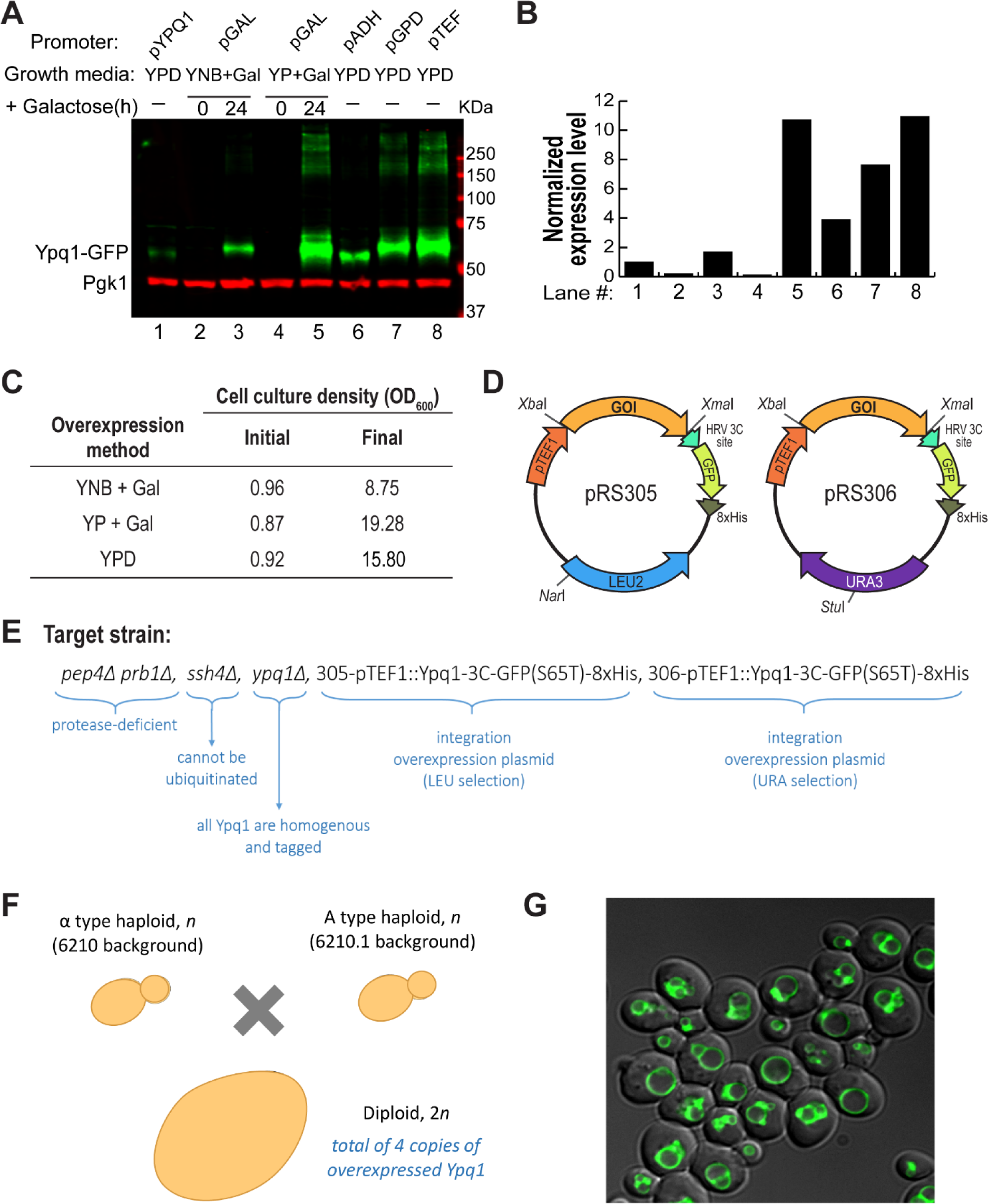
A constitutive overexpression system for lysosome/vacuole membrane proteins in *S. cerevisiae*. **A)** Ypq1-GFP expression under different induction conditions. For galactose induction: Cells expressing Ypq1-GFP-8xHis were de-repressed in media lacking leucine and supplemented with lactate or raffinose (0h +Galactose), then transferred to either YNB or YP and induced with galactose for 24 hrs (24h +Galactose). For constitutive expression: Cells expressing Ypq1-GFP-8xHis under ADH1, GPD1, or TEF1 promoters were grown in YPD, then transferred to fresh YPD and grown for 24 hrs. Ypq1-GFP expressed under its native promoter was included as a control. Each lane has lysate from 1 OD cells. Phosphoglycerate kinase (Pgk1) is used as a loading control. **B)** Quantification of (A) using GFP expression normalized to Pgk1 signal. **C)** Cell density in different induction conditions. Cells grown in YNB –leucine +lactate, then switched to YNB –leucine +galactose (YNB+Gal) vs. cells grown in YNB –leucine +raffinose, then switched to YP +galactose (YP+Gal) vs. constitutively expressing cells grown in YPD. **D)** Maps of integration plasmids expressing Ypq1-GFP-8xHis under the TEF1 promoter. Plasmids contain an HRV 3C site, and *Xba*I/*Xma*I restriction sites to clone any gene of interest. pRS305 and pRS306 are linearized with *Nar*I or *Stu*I to integrate into the yeast genome. Selections are leucine and uracil, respectively. **E)** Genetic features of the haploid overexpression yeast strain improve Ypq1-GFP-8xHis yield and stability. Shown as well are two copies of the overexpressed fusion protein. **F)** α and A haplotype haploid yeast version of (E) can be mated to produce diploid cells with 4 copies of the overexpressed fusion protein. **G)** Microscopy image of the diploid cell shows proper localization of Ypq1-GFP-8xHis on the vacuole membrane.

Galactose-inducible cells were grown in minimal media (YNB) lacking leucine and supplemented with 2% lactate or raffinose, then induced in either YNB+Galactose or rich media (YP)+Galactose for 24 h. All throughout, cells were maintained in low glucose (0.1-0.2%) to de-repress the GAL1 promoter. Meanwhile, constitutively expressing cells were grown in YP+Glucose (YPD) for 24 h. We then measured Ypq1-GFP expression in 1 OD cells (Fig. 1A-B). Although the pDDGFP2-LEU2d has been an effective expression system for many membrane proteins (Parker and Newstead, 2014), we found that, compared to endogenous levels, Ypq1-GFP expression only increased two-fold in cells induced in YNB+Galactose, and ∼11-fold in cells induced in YP+Galactose. Constitutive overexpression under pADH1 increased the expression to ∼4-fold, followed by pGPD1 to ∼8-fold and pTEF1 to ∼11-fold.

Aside from promoter strength, the number of cells (i.e., culture density) actively expressing the protein also contributes to the overall batch expression level. To test the effect of different media on culture density, we grew starter cultures to about the same culture density (optical density, OD_600_∼0.9), then switched to YNB-Leu+Gal, YP+Gal, or YPD for 24 h and measured the final OD_600_ (Fig. 1C). As expected, the lack of nutrients in the minimal medium (YNB-Leu+Gal) limited cell growth, whereas rich media YP+Gal or YPD produced very dense cultures. The galactose-inducible pDDGFP2-LEU2d plasmid relies on the selection pressure imparted by – leucine growth conditions (Parker and Newstead, 2014). Thus, cells must be grown in minimal media, which lacks many nutrients present in rich media; and under low glucose (0.1-0.2%) to keep the GAL promoter de-repressed. Together, these conditions limit cell division before and during induction. Thus, although each cell can retain high copies of the plasmid (Parker and Newstead, 2014), a low number of cells limits total Ypq1-GFP expression.

In both promoter strength and culture density tests, galactose induction via pDDGFP2-Leu2d in YP+Gal and constitutive overexpression under pTEF1 in YPD showed the highest potential for Ypq1 overexpression. In terms of ease of culture, the de-repression step in the galactose induction system requires large amounts of lactate, raffinose, and galactose, as well as prolongs culture time by up to 24 h. Thus, considering overexpression levels, cost, and technical ease, we decided to overexpress Ypq1 under a constitutive TEF1 promoter in YPD.

### A diploid overexpression strain for Ypq1 purification

We constructed pRS305 (leucine selection) and pRS306 (uracil selection) integration plasmids containing pTEF1::Ypq1-GFP-8xHis with an HRV 3C protease site at the C-terminus of Ypq1 for cleavage of the GFP-8xHis tag (Fig. 1D). Stable integration was done by linearizing the pRS305 and pRS306 plasmids with NarI or StuI, respectively, and transforming to yeast where the plasmids recombine into the genome. These two plasmids stably integrate into two different genomic loci (i.e., LEU2 and URA3).

To improve the stability of Ypq1 during purification, we built a yeast strain with several gene deletions (Fig. 1E). We deleted the major vacuolar proteases PEP4 and PRB1 to minimize proteolytic cleavage during cell lysis and membrane preparation (Wurmser and Emr, 1998). Eliminating protein ubiquitination and degradation can also improve yield. To this end, we deleted *SSH4*, an E3 ligase adaptor that recognizes Ypq1 for ubiquitination (Arines et al., 2021; Arines and Li, 2022; Li et al., 2015b). For purifying other vacuolar membrane proteins, we recommend using a triple deletion strain (*ssh4*Δ *tul*Δ *pib1*Δ), where all known vacuole membrane E3 ligase systems (Ssh4-Rsp5, DSC complex, Pib1) are disrupted (Yang et al., 2020). For membrane proteins on a different organelle, their E3 ligase systems may need to be genetically disrupted (Sardana and Emr, 2021; Sun and Brodsky, 2019). Lastly, we deleted the endogenous YPQ1 gene to ensure that all Ypq1 proteins are tagged and homogenous – the latter is critical when expressing proteins that form oligomers.

With these modifications, we generated a protease- and ubiquitination-deficient yeast strain that expressed two copies of overexpressed Ypq1-GFP-8xHis (Fig. 1E). To further increase the expression, we mated MATα (SEY6210) and MATA (SEY6210.1) haplotypes of this strain to generate a diploid strain (FAY146) expressing four copies of overexpressed protein (Fig. 1F). Using microscopy, we confirmed that the fusion protein localized properly to the vacuole membrane (Fig. 1G).

### Purification of Ypq1

Using this system, we grew a 4-L YPD culture of FAY146 for 20-24 hours (Fig. 2A). We then lysed the cells by digesting the cell wall with zymolase followed by Dounce homogenization, which we determined to improve the yield and solubility of the protein over other lysis methods such as high-pressure cell disruption (Microfluidizer^TM^) and bead-beating (Fig. S1A). We then solubilized membrane fractions in 2% DDM, as determined through our detergent screen (Fig. S1B-C), prior to affinity chromatography of the 8xHis tag onto TALON cobalt resin. Finally, Ypq1 was released using HRV 3C protease and further purified by size exclusion chromatography. Proteins from representative fractions were visualized by Coomassie staining in Fig. 2B. This purification scheme resulted in a single peak eluting at ∼13 mL fraction volume (Fig. 2C) and a ∼34 kDa band corresponding to the Ypq1 protein (Fig. 2D).

**Fig. 2.**
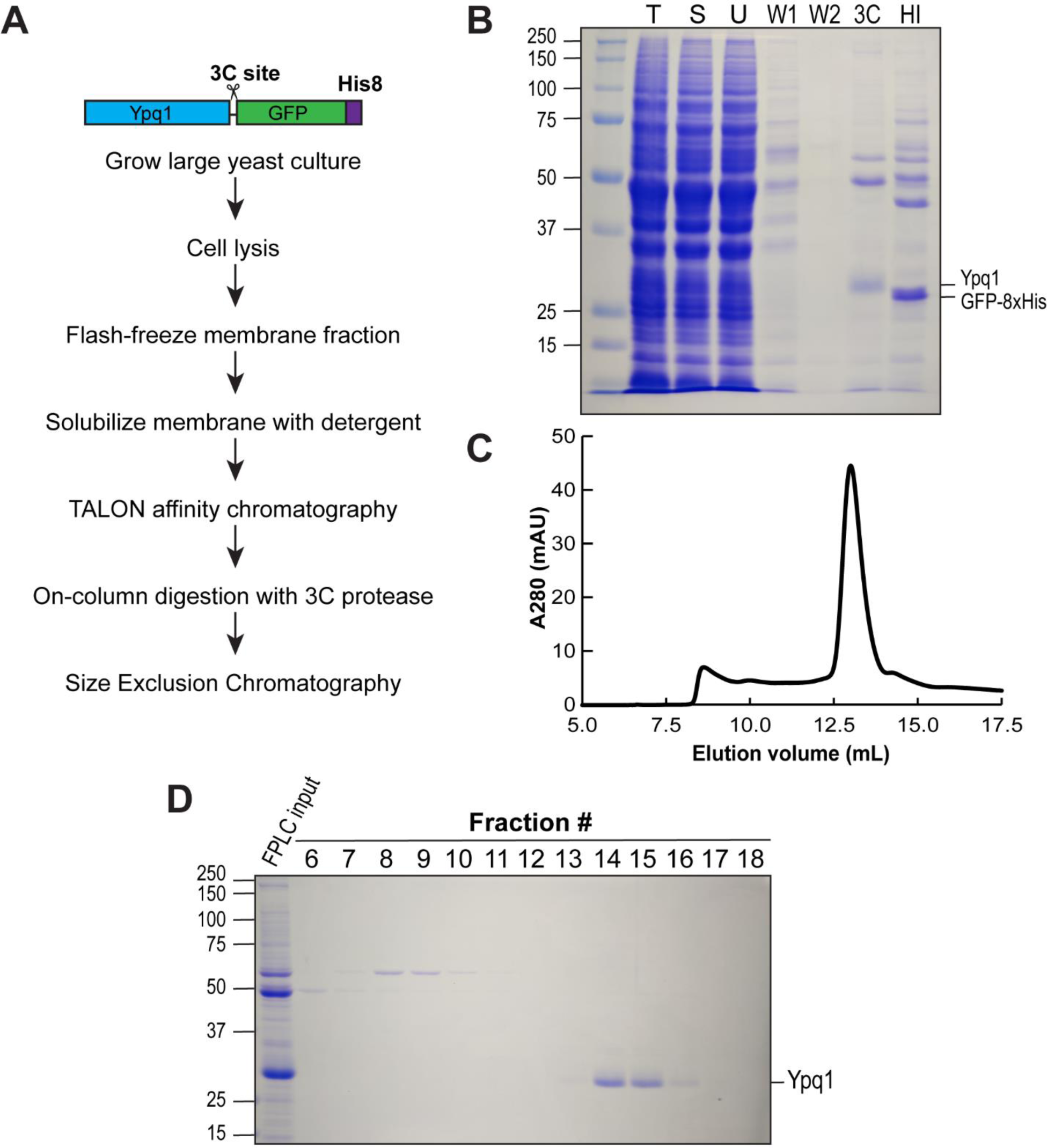
Purification of Ypq1. A) Purification scheme for Ypq1. B) Coomassie Brilliant Blue staining of purification fractions. T-Total Membrane; S - Soluble fraction; U - fraction Unbound to TALON resin; W1, W2 – TALON resin wash; 3C – eluate after 3C digestion; HI – eluate after High Imidazole wash. C) FPLC profile from a 2-L purification. D) Coomassie Brilliant Blue staining of 0.5-mL SEC fractions from (C).

During optimization, we noticed a prominent 50-kDa truncation product after cell lysis, regardless of the lysis method used (Fig. S1). The truncation product disappeared when we replaced yEGFP with GFP^S65T^ (Fig. S1D), a variant commonly used in yeast studies (Heim and Tsien, 1996). Interestingly, the two variants differ only in 3 amino acids (yEGFP→GFP^S65T^: G65T, A72S, Q80R), but somehow this has a significant impact on the stability of the fusion protein.

### Lysine stabilizes Ypq1 *in vivo* and *in vitro*

*In vivo,* Ypq1 localizes on the vacuole membrane, and its protein stability responds to cellular lysine levels (Fig. 3A). When cells are starved of lysine, Ypq1 is recognized by Ssh4, which recruits the E3 ubiquitin ligase Rsp5 (Arines and Li, 2022; Li et al., 2015b). Ubiquitinated Ypq1 then triggers the recruitment of the ESCRT machinery, which invaginates Ypq1 into the vacuole lumen where it is proteolytically degraded. In Fig. 3B, we show the subcellular localization of Ypq1-GFP before and after lysine starvation.

**Fig. 3.**
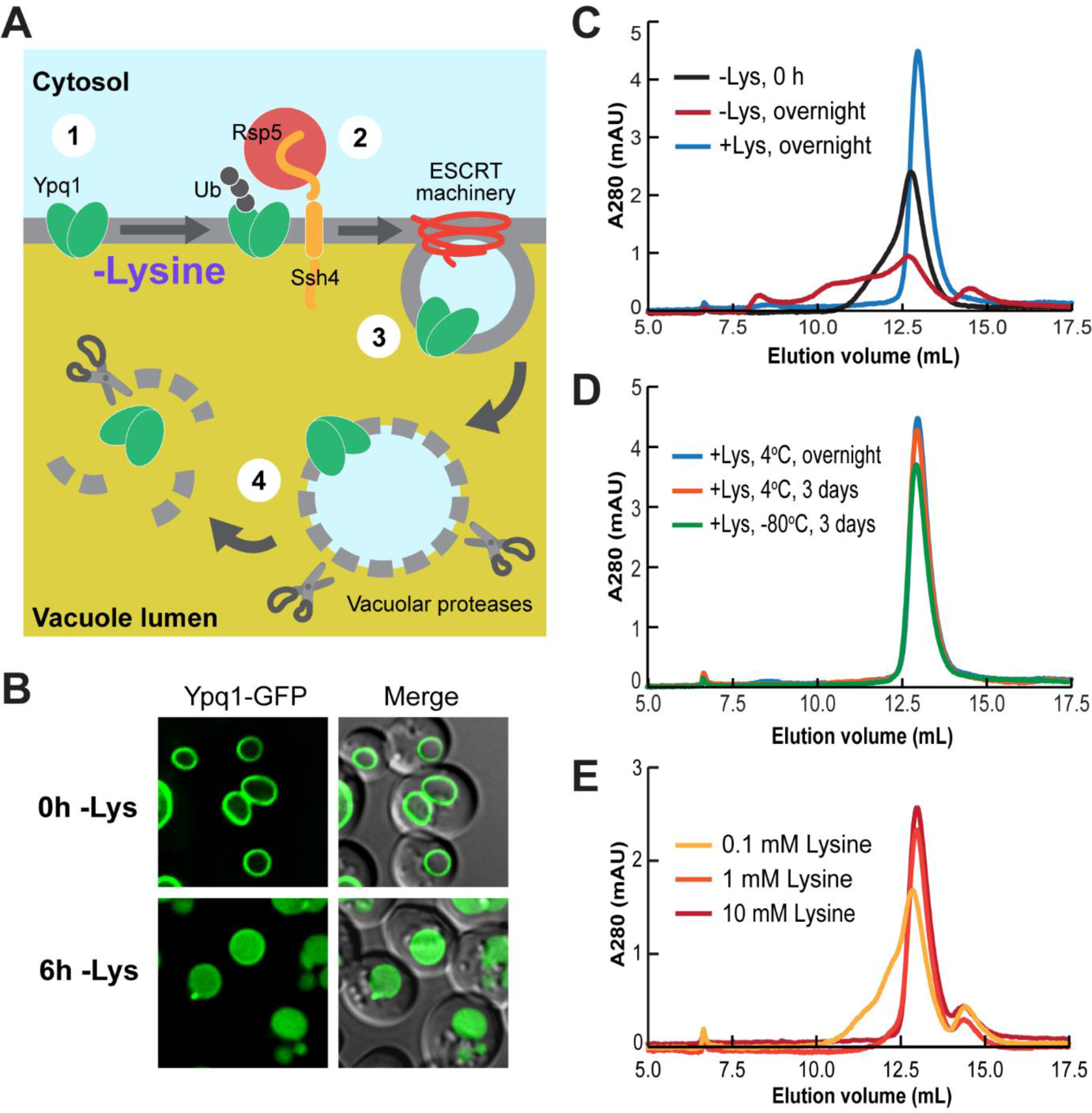
Ypq1 is stabilized by lysine *in vivo* and *in vitro*. A) Cartoon model of Ypq1 substrate-dependent degradation. 1) In the presence of lysine, Ypq1 localizes on the vacuole membrane, 2) upon lysine removal, the E3 ubiquitin ligase adaptor Ssh4 recognizes Ypq1 and recruits the E3 ligase Rsp5, which ubiquitinates Ypq1, 3) the ESCRT machinery is recruited to the membrane and internalizes Ypq1 into the vacuole lumen, 4) the internalized vesicle and Ypq1 are degraded by vacuolar proteases and lipases. B) Subcellular localization of Ypq1-GFP before or after lysine starvation. C) FPLC profiles of Ypq1 incubated in buffer with or without lysine. D) FPLC profiles of Ypq1 incubated in buffer with lysine at different storage conditions. E) FPLC profiles of Ypq1 incubated overnight in buffers with increasing lysine concentrations.

We hypothesized that purified Ypq1 is unstable without its substrate. To test this, we took equivalent amounts of Ypq1 purified under +lysine conditions, then incubated it in a buffer with or without lysine (Fig. 3C). Without lysine, the Ypq1 peak immediately shifts to the left and forms a shoulder, corresponding to the formation of higher molecular weight (MW) species, which is indicative of protein aggregation. After overnight incubation without lysine, the main peak was further reduced, and more higher-MW species appeared. This is in stark contrast with Ypq1 kept in lysine overnight, which maintains its monodispersity and elutes at ∼13 mL, similar to the initial purification (Fig. 2C). Ypq1 is stable even at 3 days +lysine and survives storage at −80°C and freeze-thaw (Fig. 3D).

To determine the minimal concentration needed to stabilize Ypq1, we tested a ten-fold concentration range for lysine based on the reported lysine pools inside the vacuole (∼5-10 mM) and the cytoplasm (∼0.5-0.8 mM)(Messenguy et al., 1980). We observed that Ypq1 is stable in 10 mM and 1 mM, but not in 0.1 mM lysine (Fig. 3E), suggesting that a specific physiological concentration of lysine is necessary to stabilize Ypq1.

Together, these results establish that lysine is required for Ypq1 stability *in vitro,* which is consistent with our prior *in vivo* observation that loss of lysine triggers Ypq1 degradation through vacuolar proteolysis. In purified conditions, Ypq1 tends to aggregate at low concentrations of lysine or upon lysine removal.

### Ypq1 is a lysine transporter

Ypq1 has been reported as a basic amino transporter (Jezegou et al., 2012; Sekito et al., 2014). To test its activity in a purified system, we reconstituted Ypq1 into yeast polar lipid extract (Avanti Polar Lipids) in the presence of 10 mM lysine to ensure protein stability. The resulting proteoliposome has a neutral internal buffer (pH 7.2) and contains 10 mM cold lysine. This concentration mimics the high lysine content in the vacuole lumen (Li and Kane, 2009; Messenguy et al., 1980). We then exchanged the proteoliposome into an external buffer containing 2 μM ^14^C-lysine (Fig. 4A). Under these conditions, if the protein is capable of lysine transport, we expect the exchange of cold and radiolabelled lysine, with the uptake of radiolabelled lysine into the liposomes driven by the outward-directed lysine gradient. At 10 mins, we saw an accumulation of ^14^C-lysine inside proteoliposomes, in contrast with liposomes without Ypq1. The ^14^C-lysine uptake was reduced upon the addition of external 10 mM cold lysine, which not only disrupts the lysine gradient but also outcompetes radiolabelled lysine for the substrate binding site. Thus, we confirm that Ypq1 transports lysine across the lipid bilayer *in vitro*.

**Fig. 4.**
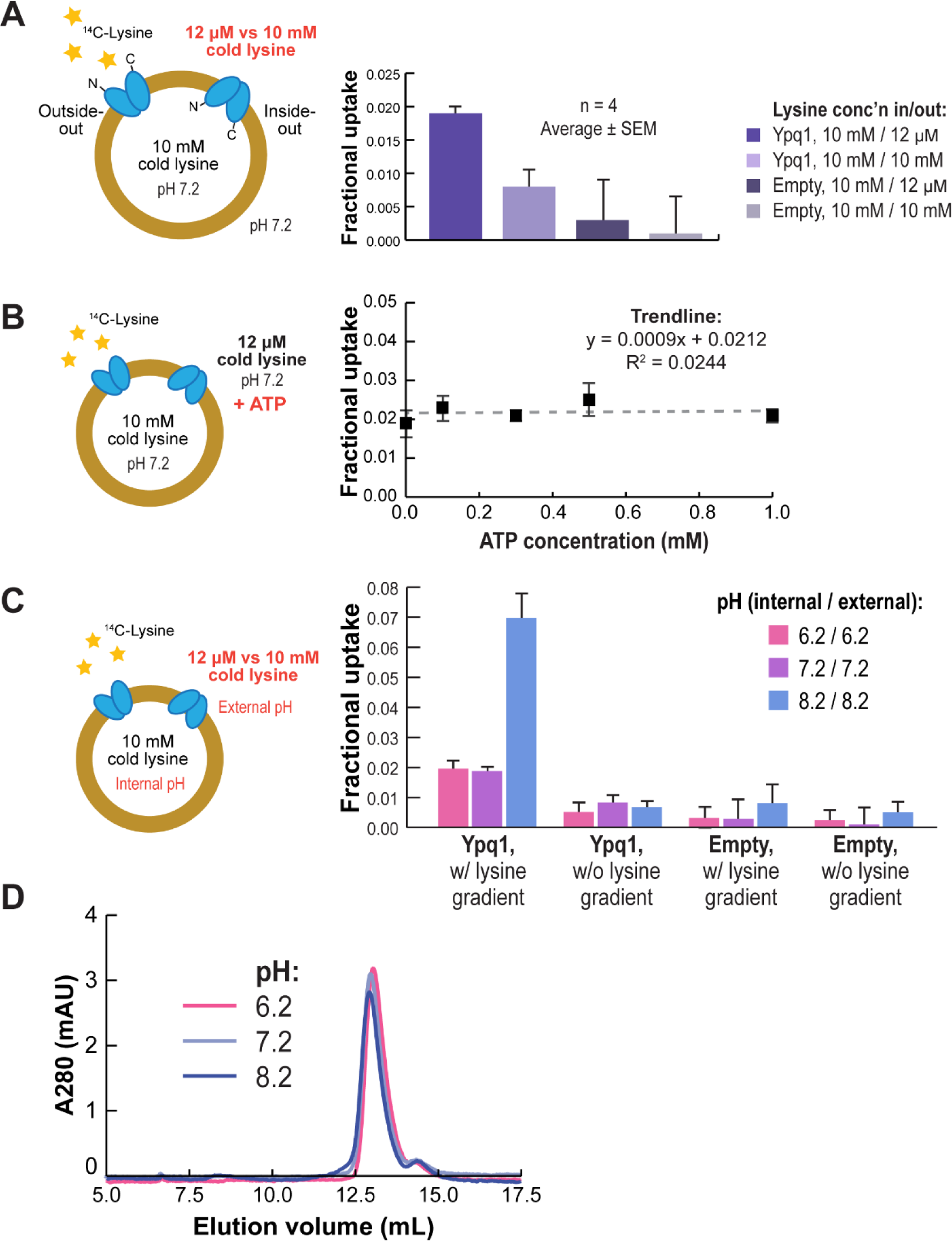
Ypq1 is a lysine transporter. For each panel, Ypq1 was reconstituted into Yeast Extract Polar liposomes (4 ug protein/mg lipid) with the illustrated intra- and extra-liposomal conditions. Reactions were started by adding 2 uM ^14^C-lysine to the extra liposomal buffer and incubating for 10 mins. Fractional uptake values are the mean and SEM of 4 independent experiments using protein from 2 independent protein preps. Empty liposomes do not contain any protein and represent basal uptake or any leakiness introduced by the buffer conditions. A) Transport activity of Ypq1 in the presence or absence of an outward lysine gradient. B) Transport activity of Ypq1 in response to ATP. Proteoliposomes were preincubated with different concentrations of ATP for 30 s before adding ^14^C-lysine and incubating for 10 mins. The trendline denotes that ATP does not affect Ypq1 transport activity. C) Transport activity of Ypq1 in different pH conditions and in the presence or absence of an outward lysine gradient. D) FPLC profiles showing the stability of Ypq1 incubated overnight in different pH.

It has been shown that ATP is important in the transport activity of Ypq1 in vacuolar membrane vesicles obtained by density gradient centrifugation (Sekito et al., 2014). Sekito and colleagues have shown that adding the proton ionophore CCCP abolishes ATP-dependent transport. Thus, they propose that ATP is a substrate for v-ATPase, which generates the H^+^ gradient that drives the Ypq1 function (Sekito et al., 2014).

To test this model, we pre-incubated Ypq1 proteoliposomes in increasing concentrations of ATP for 30 s, then started the reaction by adding 2 μM ^14^C-lysine (Fig. 4B). After 10 mins of incubation, we measured ^14^C-lysine uptake and found no significant increase in uptake even at 1 mM ATP, supporting the previously proposed model that lysine transport function by Ypq1 does not directly depend on ATP.

### pH-dependence of lysine transport by Ypq1

To test if pH influences Ypq1 function, we measured ^14^C-lysine uptake in buffers at pH 6.2, 7.2, and 8.2 (Fig. 4C). We were expecting that uptake would increase in acidic pH, consistent with the acidic pH of the vacuole lumen (pH 5-6.5, Li and Kane, 2009). However, ^14^C-lysine uptake at pH 6.2 was about the same as the uptake at pH 7.2, in both the presence or absence of an outward-directed cold lysine gradient. Instead, uptake increased by more than 3-fold at pH 8.2, suggesting that a basic pH on at least one side of the membrane stimulates Ypq1 function. We confirmed that the differences in uptake are not due to Ypq1 instability because Ypq1 in detergent micelles was stable at all pH conditions, provided that 10 mM lysine is maintained (Fig. 4D).

We next sought to test whether a pH gradient could drive lysine uptake. However, we were unable to reduce the intraliposomal 10 mM lysine without protein instability. Therefore, we analyzed radiolabelled lysine uptake driven by both lysine and proton gradients, compared to substrate uptake driven by the lysine gradient alone. Relying only on an outward-directed cold lysine gradient, a symmetric internal and external pH of 6.2 resulted in minimal ^14^C-lysine uptake (Fig. 5A, blue trace). The equilibrium uptake increased substantially when we applied a 100-fold H^+^ gradient (pH 6.2 internal/pH 8.2 external) (Fig. 5A, yellow trace). The combined effect of the outward-directed cold lysine gradient together with the H^+^ gradient drove a larger fractional uptake than the lysine gradient alone, with symmetric internal/external pH of 8.2 (Fig. 5B). This suggests that a large, outward-directed H^+^ gradient across the lipid bilayer also drives uptake of radiolabelled lysine, with an additive effect on the equilibrium uptake. Application of the opposite pH gradient (internal/external pH 8.2/6.2) abolished uptake of radiolabelled lysine (Fig. 5C), further suggesting that in these experiments, the proton gradient acts as a driving force rather than as a biochemical modulator of protein activity.

**Fig. 5.**
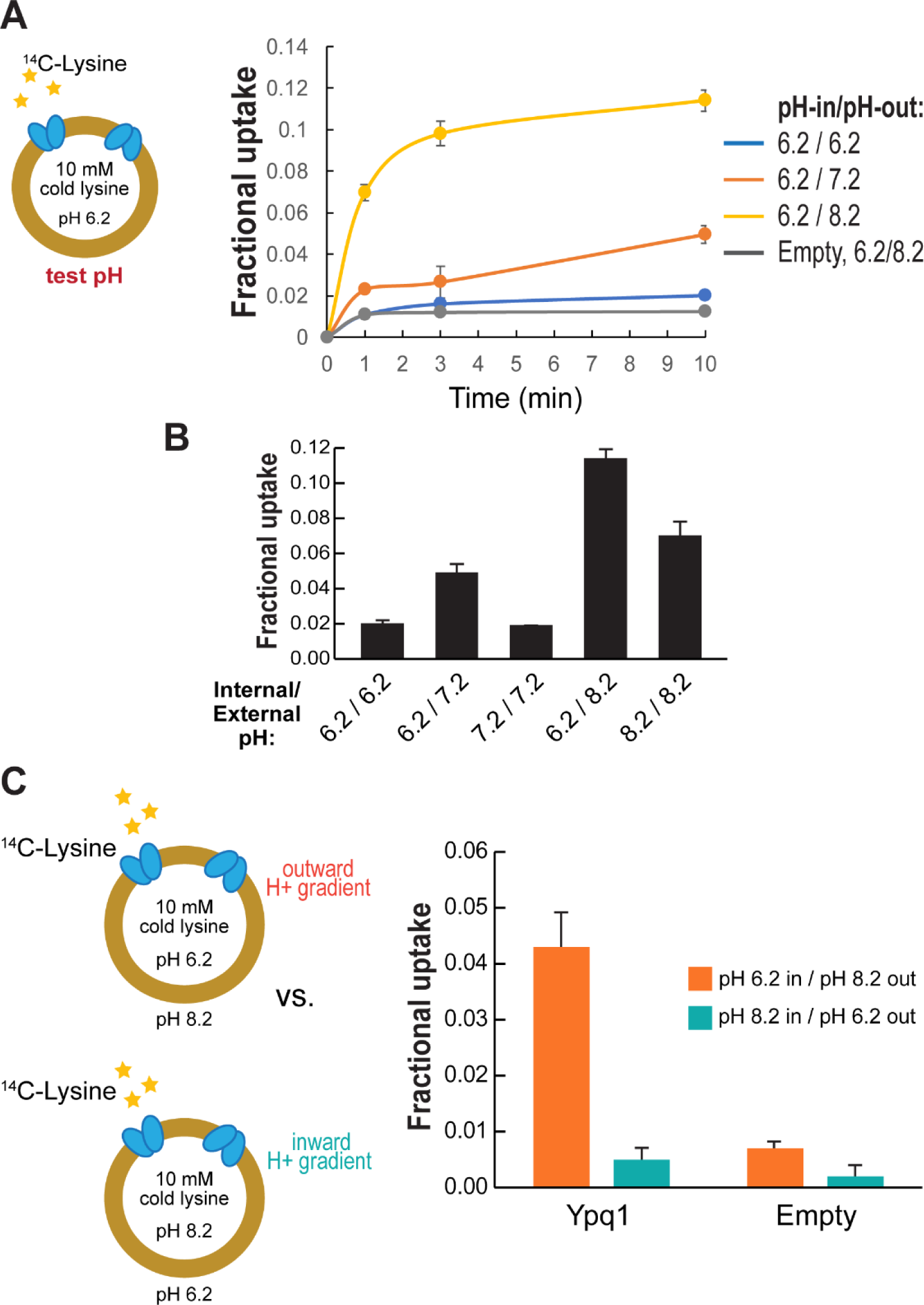
Ypq1 is likely a proton-coupled lysine antiporter. Time course of ^14^C-lysine uptake into proteoliposomes with the indicated internal and external pH. In all samples, the intraliposomal buffer contains 10 mM lysine while the external buffer does not have lysine. Transport values are the mean ± SEM of 4 independent experiments using protein from 2 independent protein preps. A) ^14^C-lysine uptake of Ypq1 proteoliposomes after 10 min in the presence of the indicated liposomal pH gradients. B) Comparison of 10-min fractional uptake in liposomes with symmetric pH vs. liposomes with a pH gradient. C) ^14^C-lysine uptake activity of Ypq1 proteoliposomes in the presence of an outward vs. inward H^+^ gradient.

### Ypq1 stability and transport are differentially affected by lysine and arginine

Previous biochemical studies have reported that Ypq1 could also transport arginine, but to a lesser extent compared to lysine (Cools et al., 2020; Kawano-Kawada et al., 2019; Sekito et al., 2014). Some earlier studies have even hinted that Ypq1 could be transporting basic amino acids in general, i.e., lysine, arginine, and histidine (Jezegou et al., 2012). To test if there is a link between basic amino acid transport and Ypq1 downregulation, we expressed Ypq1-GFP in yeast that is auxotrophic to lysine, arginine, histidine, and leucine. We first grew the cells in rich media and then starved them in media lacking basic amino acids (Fig. 6A). As a control, we incubated cells in rich media, which contains all amino acids, and media lacking the non-charged amino acid leucine. As expected, Ypq1-GFP was internalized into the vacuole lumen after lysine starvation. However, no degradation was detected in –arginine, –histidine, and –leucine.

**Fig. 6.**
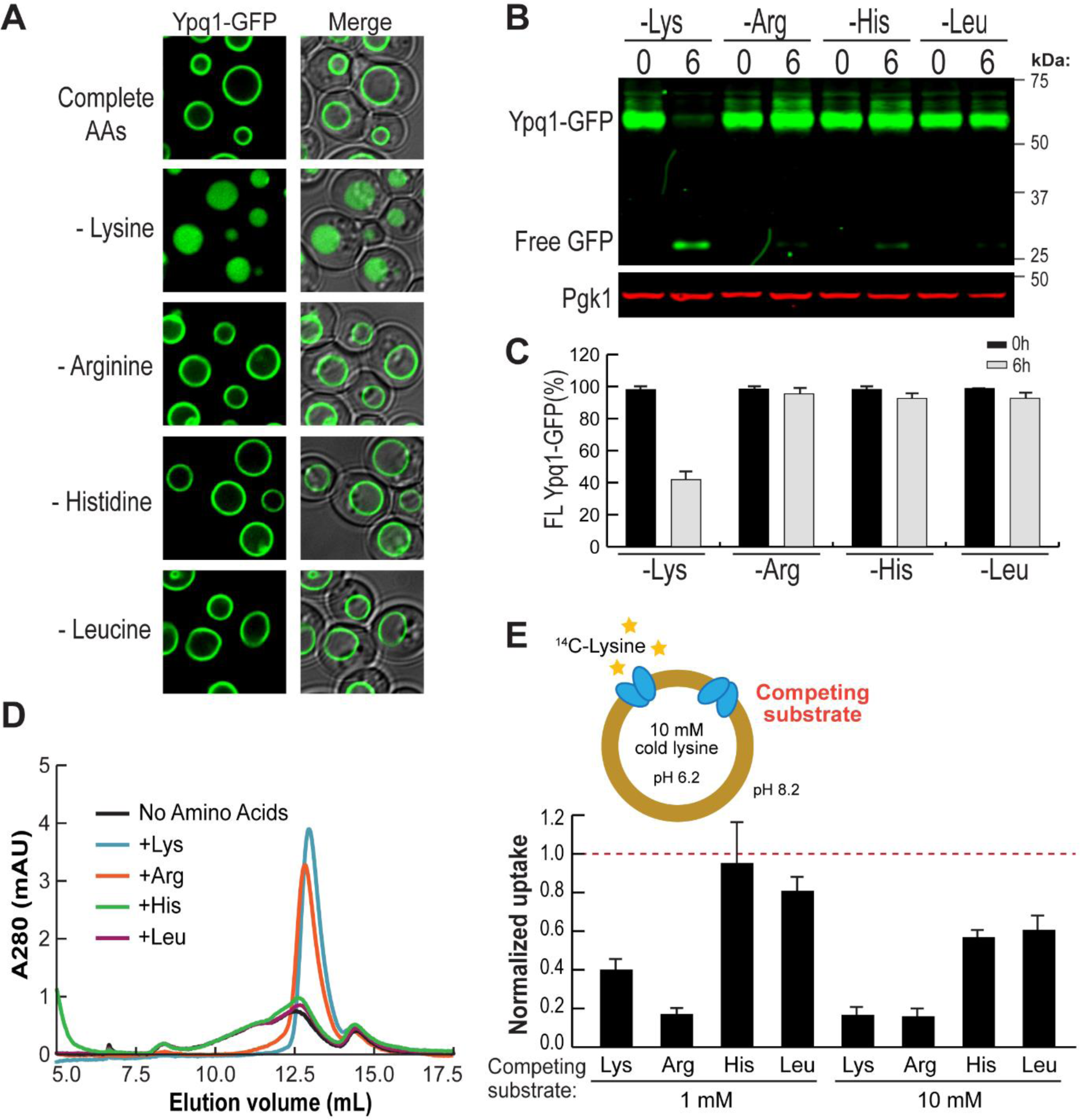
Ypq1 stability and transport are affected by the presence of lysine and arginine. A) Subcellular localization of Ypq1-GFP after growing cells for 6 h in the presence of rich media or in media lacking the designated amino acid. B) Western blot analysis of Ypq1-GFP degradation at 0h or 6h in the absence of different amino acids. Upon vacuolar degradation of Ypq1-GFP, the full-length band (64 kDa) signal decreases while the free GFP band (27 kDa) increases. C) Quantification of protein levels was calculated as full-length Ypq1-GFP divided by total GFP signal (i.e. full-length Ypq1-GFP + free GFP). Error bars represent SD (n = 3). D) FPLC profiles showing the stability of Ypq1 incubated overnight in different amino acids. Substrate selectivity of Ypq1 was tested by incubating proteoliposomes preloaded with 10 mM lysine (pH 6.2) with an external buffer (pH 8.2) containing 1 mM or 10 mM of the indicated amino acids. ^14^C-lysine uptake was measured after 3 mins and was normalized to the uptake of proteoliposomes incubated in external buffer without amino acids. Transport values are the mean ± SEM of 4 independent experiments using protein from 2 independent protein preps.

We demonstrated this further by Western blot, wherein vacuolar degradation is characterized by the loss of the full-length GFP fusion protein and the accumulation of a free GFP band (Fig. 6B-C). Consistent with imaging data, Ypq1-GFP was degraded in –lysine but not in other media. However, we did observe a very faint free GFP band in –arginine and –histidine but not in –leucine. This suggests that *in vivo*, Ypq1 degradative regulation responds mostly to lysine starvation and perhaps to arginine and histidine starvation, but only to a very minimal extent.

We further tested the effect of these amino acids on the stability of Ypq1 in detergent micelles(Fig. 6D). We incubated purified Ypq1 in a buffer containing the test amino acids (10 mM) overnight and checked for peak monodispersity via FPLC. Similar to lysine, arginine also stabilized Ypq1, but caused a slight leftward shift of the peak compared to lysine. In contrast, histidine did not stabilize Ypq1, and the FPLC profile looked no different from the FPLC profiles in leucine buffer and the no amino acid control.

To determine whether other amino acids are competitive with lysine binding by Ypq1, we reconstituted Ypq1 in proteoliposomes preloaded with an acidic buffer (pH 6.2) and 10 mM cold lysine (Fig. 6E). We then incubated these into an external buffer (pH 8.2) containing radiolabelled lysine and cold test amino acids for 3 mins (instead of 10 mins to avoid saturation). At a low concentration (1 mM), lysine and arginine, but not histidine or leucine, reduced ^14^C-lysine accumulation in the proteoliposome. Unexpectedly, we noticed a stronger effect by arginine, which did not further increase at 10 mM. At a high concentration (10 mM), cold lysine further competed with ^14^C-lysine and reduced its uptake at a similar level as 10 mM arginine. Notably, 10 mM histidine and 10 mM leucine only slightly inhibited ^14^C-lysine uptake.

Together, these results show that basic amino acids have a differential effect on Ypq1 stability and transport. *In vivo*, Ypq1 ubiquitination and degradation respond mostly to lysine withdrawal, but not to other basic amino acids. *In vitro*, Ypq1 is stabilized by both lysine and arginine, but not histidine. It is possible that lysine and arginine are metabolized at different rates *in vivo* after being removed from the media.

### The stability of Ypq1 degradation mutants is consistent *in vivo* and *in vitro*

Previously, we identified two sets of Ypq1 mutants with a defect in substrate-dependent degradation (Arines et al., 2021). One set resists degradation even after lysine starvation, while another set is constitutively degraded even in the presence of lysine. We asked if these mutants have different stabilities *in vitro,* and so influence their regulation. To this end, we purified two representative mutants, P229S,Q230R (herein referred to as PQmut) and M73D, and tested their stability in detergent micelles.

A hallmark of the PQ-loop protein family, which includes the vacuolar basic amino acid transporters Ypq1-3 in yeast (Arines et al., 2021; Cools et al., 2020; Kawano-Kawada et al., 2019; Manabe et al., 2016; Sekito et al., 2014); the lysine/arginine transporter LAAT-1 in *C. elegans* (Liu et al., 2012); the mammalian basic amino acid transporter PQLC2 (Leray et al., 2021); the KDEL receptor (Brauer et al., 2019); and the sugar transporters SemiSWEET (Latorraca et al., 2017; Lee et al., 2015) and SWEETs (Han et al., 2017; Tao et al., 2015), is the presence of two conserved proline-glutamine (PQ) dipeptide motifs. Based on structural studies, the PQ motif acts as a molecular hinge that allows PQ-loop transporters to change conformations during transport. In our *in vivo* degradation studies, we found that mutating the second PQ motif (P_229_Q_230_) results in the loss of the recognizability by Ssh4 and blocks Ypq1 degradation. The second mutant, M73D, is constitutively recognized by Ssh4 and degraded quickly (Arines et al., 2021).

In Fig. 7A, we show that PQmut and M73D escape the normal regulatory response – PQmut is not degraded and stays on the vacuole membrane even in –lysine media, while M73D is degraded in the vacuole lumen even in +lysine media. We know that M73D still trafficks properly and that its recognition only happens on the vacuole membrane, because when SSH4 is deleted, the M73D mutant localizes correctly on the vacuole membrane (Fig. 7B).

**Fig. 7.**
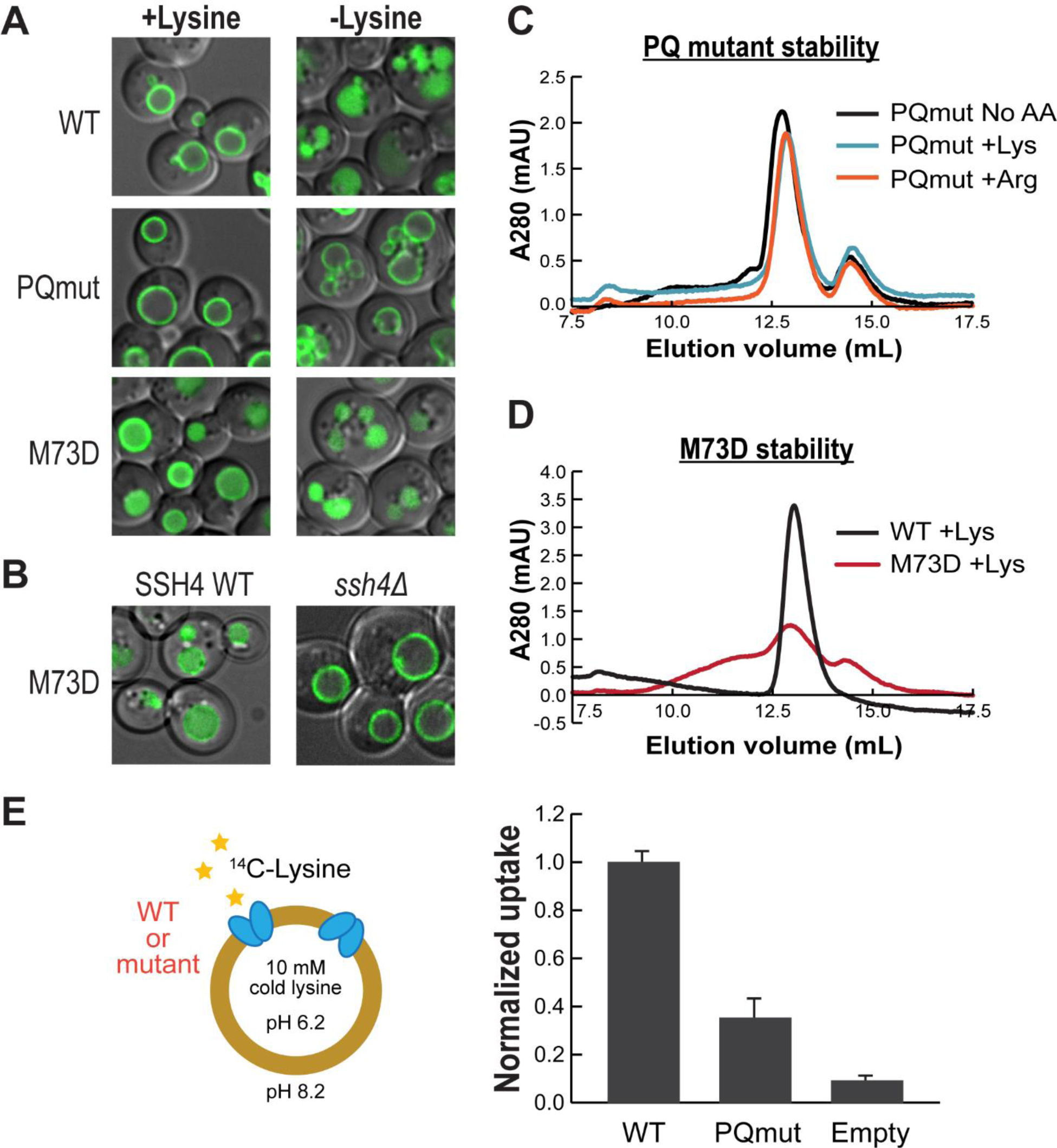
Stability and transport of Ypq1 degradation mutants. A) Subcellular localization of wild-type (WT) vs. degradation-blocking (PQmut) and constitutively-degrading (M73D) mutants of Ypq1-GFP. B) Subcellular localization of Ypq1(M73D)-GFP in the presence or absence of Ssh4. C) FPLC profiles showing the stability of purified Ypq1(PQmut) incubated overnight in buffer +lysine, +arginine, nor no amino acids (AA). D) FPLC profiles showing the stability of purified WT Ypq1 or Ypq1(M73D) in the presence of lysine. E) Normalized ^14^C-lysine uptake activity of WT Ypq1 vs Ypq1(PQmut). Proteoliposomes preloaded with pH 6.2 buffer containing 10 mM lysine were exchanged into an external pH 8.2 buffer. ^14^C-lysine uptake was measured after 10 mins and was normalized to the uptake of WT Ypq1. Transport values are the mean ± SEM of 4 independent experiments from two independent protein preps.

Consistent with its stability *in vivo*, purified PQmut is stable in all buffers tested, even when amino acids are absent (Fig. 7C). We had difficulties purifying M73D, mainly due to its low yield and propensity to aggregate. With the limited M73D mutant we managed to purify, we tested its stability in buffer +lysine and saw a low and broad profile, unlike the monodisperse wild-type peak (Fig. 7D). Thus, both mutants showed consistent stabilities *in vivo* and *in vitro*, suggesting that inherent protein stability influences the recruitment of the ubiquitination machinery.

We also asked whether these mutants have a defect in transport by reconstituting the mutants into proteoliposomes applied with a pH gradient (pH 6.2 in/pH 8.2 out) and preloaded with 10 mM lysine (Fig. 7E). We then measured the uptake of ^14^C-lysine after 10 mins incubation. Similar to previous studies on other PQ-loop proteins (Cools et al., 2020; Kawano-Kawada et al., 2019; Liu et al., 2012), mutating the PQ motif reduced uptake to ∼40%, consistent with its importance in transport. Due to the propensity to aggregation and low yield, we could not get enough M73D protein for transport experiments.

## DISCUSSION

### Lessons from Ypq1 purification

In this study, we developed a method to overexpress, purify, and reconstitute vacuole membrane proteins into proteoliposomes, and used this method to interrogate Ypq1 stability and transporter activity. The lessons we learned during the development of the method could be emulated for other membrane proteins.

First, constitutive overexpression in rich media and an increase in the copy number (up to 4 copies) of overexpression constructs gave the best yield for Ypq1. Second, our purifications emphasized a biochemical feature of many transporter proteins, that purification in the presence of physiological concentrations of substrate improves yield and stability. We propose that there may be a link to the *in vivo* regulation of protein stability by substrate concentrations (Keener and Babst, 2013; Li et al., 2015a; Lin et al., 2008; Sardana et al., 2019). Third, disrupting the ubiquitination machinery by deleting Ssh4 improved the yield. For vacuole membrane proteins with unknown E3 ligases, an *ssh4Δ tul1Δ pib1Δ* strain can be used instead. Lastly, switching the tag from yEGFP to GFP(S65T) eliminated Ypq1 truncation, but the reason is not yet apparent.

### A pH-gated mechanism of lysine transport

Here, we demonstrated that Ypq1 is a lysine transporter that does not directly use ATP for transport. Furthermore, we found that Ypq1 transport is activated at basic pH, and that uptake into proteoliposomes is further increased upon applying an outward-directed pH gradient, consistent with a model where proton export from proteoliposomes drives lysine uptake. We interpret these results to mean that Ypq1 catalyzes proton/lysine antiport. In the cellular context, this would imply that the acidic pH in the vacuole lumen drives lysine uptake into the vacuole, consistent with a metabolic role of storing and sequestering this amino acid under lysine-replete conditions (Li and

Kane, 2009; Messenguy et al., 1980). Although we cannot rule out other possible models, such as an interplay between substrate concentration and pH that modulates biochemical activity, our proposal that Ypq1 acts as a proton-coupled antiport provides a plausible model for the biochemical and physiological data obtained so far.

We also found that, aside from serving as a driving force, a basic pH on at least one side of the vacuolar membrane is necessary for protein activity. Because we do not know the direction of protein insertion in the liposomes, our experiments do not directly probe whether the luminal or cytosolic side is responsible for this pH dependency. However, given the low pH within the yeast vacuole, we assume that this pH regulation occurs on the cytosolic side of the protein. Although pH 8.2 is higher than the yeast cytoplasm, many metabolic enzymes exhibit pH optima above the physiological pH (Luzia et al., 2022; Simcikova and Heneberg, 2019). This provides a greater dynamic range for pH-dependent regulation of activity within the physiological range, suggesting that Ypq1 activity might be tuned to respond to perturbations in the cytosolic pH. Transient increases in cytosolic pH are associated with nutrient-replete conditions, providing another link between metabolic flux and lysine uptake for storage in the vacuoles.

We also showed that arginine exerts similar effects as lysine on Ypq1 stability *in vitro* and that arginine is competitive with lysine uptake. However, our assays do not distinguish arginine binding and transport, and arginine withdrawal does not trigger Ypq1 degradation in cells. Thus, whether Ypq1 plays a physiologically important role in arginine transport remains an open question.

### Coupling of *in vitro* stability and transport to *in vivo* degradation

The stability of purified Ypq1 provides new information to explain its substrate-dependent degradation. We found that the stability of wild-type, degradation-resistant PQmut, and constitutively-degrading M73D are consistent *in vivo* and *in vitro*. This provides evidence that lysine directly regulates Ypq1 folding/stability and *not* the E3 ligase Rsp5 or its adaptor Ssh4.

The nature of these changes is not yet fully understood. Previously, we proposed that transport-associated conformation changes on Ypq1 expose polar transmembrane residues that are recognized by Ssh4 (Arines et al., 2021). In our current work, M73D instability suggests that it is either misfolded or trapped in a conformation that persistently exposes these polar residues, which can lead to aggregation *in vitro*. However, it is puzzling because M73D, although very unstable *in vitro*, can surpass ER quality control and localize properly on the vacuole membrane in the absence of Ssh4 (Fig. 7B). M73D might be more stable in membranes than detergent micelles. Another possibility could be that there might be transmembrane chaperones that are shielding the exposed polar residues. These putative chaperones might be missing after Ypq1 purification.

Conversely, the PQ mutant is resistant to lysine withdrawal *in vivo* and stable *in vitro* after lysine is removed from the buffer. These consistent results suggest that the PQ mutant is properly folded, and its folding is no longer regulated by the substrate lysine. The crystal structures of the PQ family members, including SWEET and cystinosin, indicated that the proline functions as a helix breaker for the transmembrane helices. The helix breaker also provides a structural basis for conformation change and probably is necessary to transiently expose the charged residues during lysine translocation (Lee et al., 2015; Tao et al., 2015). Mutating the PQ motif could block this conformation change and reduce the lysine translocation efficiency. In line with this hypothesis, the *in vitro* lysine transportation efficiency is reduced by 65% (Fig. 7E).

Most of the lysosomal membrane proteins remain poorly studied. Establishing a platform to overexpress, purify, and reconstitute lysosomal membrane transporters opens the door to systematically characterizing their function.

## MATERIALS AND METHODS

### Construction of constitutively overexpressing strains

The overexpression plasmid was constructed in several steps to generate Ypq1-3C-GFP(S65T)-8xHis expressed under the ADH1, GPD1, or TEF1 promoters. A plasmid expressing Ypq1-3C-GFP(S65T)-8xHis under the ADH1 promoter was built by Gibson assembly and ligated into pRS305 and pRS306 vectors. The promoter region was cloned to be flanked by *Sac*I and *Xba*I restriction sites. Next, the ADH1 promoter was replaced via *Sac*I/*Xba*I restriction enzyme cloning with promoter regions of GPD1 and TEF1, as amplified via PCR from SEY6210.

The overexpression plasmids under the pRS305 and pRS306 vectors were integrated into the yeast genome by digesting with *Nar*I and *Stu*I, respectively, and transforming into α type haploid yeast deleted for PEP4, PRB1, PRC1, SSH4, and YPQ1 to generate FAY139 (6210, *SSH4Δ*::Kan^r^, *PEP4Δ*::Hyg^r^, *YPQ1Δ*::his3, *PRB1Δ*::trp1, 305-pTEF1::Ypq1-3C-GFP(S65T)-His8::tadh (LEU), 306-pTEF1::Ypq1-3C-GFP(S65T)-His8::tadh (URA)). We transformed the same plasmids into the corresponding A type haploid yeast to generate FAY141. Both FAY139 and FAY141 express two copies of pTEF1::Ypq1-3C-GFP(S65T)-8xHis, with each copy integrated into the LEU2 and URA3 loci. FAY139 and FAY141 were then mated and screened for diploid cells. The expression in these cells was confirmed by Western blot, and the cellular localization was checked by microscopy.

### Generation of mutant strains

Site-directed mutagenesis was performed on PFA390 (pRS305-pTEF1::Ypq1-3C-GFP(S65T)-8xHis) and PFA391 (pRS306-pTEF1::Ypq1-3C-GFP(S65T)-8xHis) with Novagene KOD Hot-Start DNA polymerase (71086; MilliporeSigma). Mutant plasmids were then integrated into yeast as described above.

### Spheroplasting and preparation of yeast membrane fractions

Proteins were expressed from a 4-L culture grown in YPD at 26°C for 18-24 h. Cells were divided into 1-L portions (12,000-18,000 OD cells) and harvested using a Fiberlite F10-4x1000 LEX rotor at 7,300 rpm for 12 mins. In the next steps, buffers were supplemented with 1x cOmplete™ EDTA-free Protease Inhibitor Cocktail (21169500; Roche) and 1 mM phenylmethylsulfonyl fluoride (PMSF). To each 1-L pellet, 25 mL weakening buffer (100 mM Tris-Cl (pH 8.8), 10 mM DTT) was added and incubated for 5 mins to soften the cell wall. Cells were then spun at 5,000 rpm (Sorvall SS-34 rotor) for 5 mins at 4°C and resuspended in 12.5 mL Spheroplasting buffer (2% glucose, 1x amino acids, 1 M sorbitol, 20 mM Tris-HCl, 1x YNB, pH 7.2, filtered). Cell walls were digested with zymolase 100-T (120493-1; Amsbio) at a ratio of 75 uL zymolase (5 mg/mL) per 1,000 OD cells. Digestion reactions were incubated with nutating at 30°C for 1 hr, relieving the pressure by opening the tube every 5-10 mins. The resulting spheroplasts were then spun at 7,500 rpm for 5 mins at 4°C, and resuspended in 20 mL Lysis buffer (20 mM HEPES-KOH (pH 7.2), 50 mM KOAc, 1 mM EDTA, 10% glycerol, filtered). Cells were lysed by Dounce homogenization (25 strokes) on ice. The membrane fraction was collected through a 30-min 15,000 rpm spin (Sorvall SS-34 rotor) at 4°C, and washed with 20 mL Lysis buffer without EDTA (20 mM HEPES-KOH (pH 7.2), 50 mM KOAc, 10% glycerol, filtered). The washed membrane fraction was again spun at 15,000 rpm for 30 mins at 4°C, flash-frozen in liquid nitrogen, and stored at −80°C.

### Purification of Ypq1

All steps were done at 4°C or on ice. Frozen membranes corresponding to 2-L culture fractions were thawed and resuspended in Lysis buffer supplemented with lysine (20 mM HEPES-KOH, 50 mM KOAc, 10% glycerol, 10 mM lysine, pH 7.2, filtered). Membrane fractions were resuspended by Dounce homogenization (20 strokes) on ice, and solubilized by adding DDM (D310S; Anatrace) to 2% (w/v) and nutating for 1 hr. Insoluble materials were removed by ultracentrifugation at 100,000xg (Thermo Scientific T-647.5 rotor) for 50 mins. Affinity purification of Ypq1-3C-GFP-8xHis was done by incubating the supernatant with 3 mL TALON cobalt resin (635502; TaKaRa Bio) pre-equilibrated with Column buffer (20 mM HEPES-KOH (pH 7.2), 50 mM KOAc, 5% glycerol, 0.1% DDM, 10 mM Lysine, 10 mM imidazole) for at least 1 hr in an end-over-end rotator. TALON beads were collected by centrifugation at 1000xg for 5 mins, packed into a gravity flow column (29924; Thermo Scientific), and washed with 20 column volumes of Wash buffer (20 mM HEPES-KOH (pH 7.2), 50 mM KOAc, 5% glycerol, 0.1% DDM, 10 mM lysine, 20 mM imidazole). Untagged Ypq1 was eluted by on-column digestion by incubating the resin with 200 uL HRV 3C protease (88946; Thermo Scientific) for 16 hrs. The next morning, elutions were collected, concentrated, and loaded onto a Superdex 200 Increase 10/300 GL column (28-9909-44; Cytiva). Ypq1 was further purified by gel filtration in SEC buffer (20 mM HEPES-NaOH, 100 mM NaCl, 10 mM lysine, 0.05% DDM, pH 7.2). Peak fractions were pooled and used immediately for reconstitution, or supplemented with glycerol to 10%, flash-frozen in liquid nitrogen, and stored at −80°C. Protein purity was checked by SDS-PAGE and stained with Coomassie Brilliant Blue dye. Protein identity was further confirmed by mass spectrometry (Proteomics Core Facility, Michigan State University).

### Stability testing of purified Ypq1

For testing the stability of Ypq1 in different amino acids: Pooled peak fractions were divided into equivalent volumes, loaded onto Amicon 0.5-mL 50K MWCO concentrator tubes (UFC505024; Millipore), and centrifuged at 14,000xg at 4°C to 100-uL volumes. Samples were then washed 3-5 times with 400 uL of the corresponding buffer containing the desired test amino acid, spinning at 14,000xg each time. At the last wash, samples were transferred to a new tube and stored overnight at 4°C. The next day, samples were loaded onto a Superdex 200 Increase 10/300 GL column pre-equilibrated with the test buffer. Samples were resolved via gel filtration and the resulting FPLC profiles were recorded.

For testing the stability of Ypq1 in different storage conditions: 16 ug protein from the pooled peak fractions were either stored at 4°C, or flash-frozen with 10% glycerol and stored at −80°C. Afterward, samples were loaded onto a Superdex 200 Increase 10/300 GL column pre-equilibrated with SEC buffer, and resolved via gel filtration.

### Reconstitution of Ypq1 into proteoliposomes

To prepare 1 mL of proteoliposomes (4 ug protein/mg lipid), 400 ul of Yeast Extract Polar lipids dissolved in chloroform (25 mg/mL) (190001; Avanti Polar Lipids) were dried under a slow stream of nitrogen gas and washed with pentane at least three times to remove any trace of chloroform. The lipid film was sonicated in intraliposomal buffer (20 mM HEPES, 100 mM NaCl, 10 mM lysine, at the desired pH) supplemented with 10 mM DDM, and allowed to solubilize at room temperature for at least 30 mins. Using an Amicon 0.5-mL 50K MWCO concentrator tube, freshly purified Ypq1 was buffer-exchanged by washing at least three times with SEC buffer (20 mM HEPES-NaOH, 100 mM NaCl, 10 mM lysine, 0.05% DDM) modified to the desired pH. Ypq1 equivalent to 40 ug was then combined with the solubilized lipid and incubated at 4C for 1 hour. In the meantime, Bio-Beads SM2 (152-3920; Bio-Rad) were activated with methanol, washed twice with ddH_2_O, and equilibrated in the desired intraliposomal buffer. Detergent removal was done by incubating the lipid-protein mixture in Bio-Beads at 4°C overnight in an end-over-end rotator. The next day, proteoliposomes were collected, flash-frozen in liquid nitrogen, and stored at −80°C.

### Radioactive transport assays

Radioactive transport assays were done following ^47^ with some modifications. Briefly, proteoliposomes were freeze-thawed three times by flash-freezing in liquid nitrogen and slow-thawing at room temperature. Proteoliposomes were then extruded 21 times through a 400-nm membrane filter. To exchange into the desired extraliposomal buffer, extruded proteoliposomes were passed over a Sephadex G-50 column pre-equilibrated with the desired buffer (20 mM HEPES, 200 mM sorbitol, test pH, test substrate). Transport was initiated by adding 1:100 volume of hot substrate (^14^C-lysine (NEC280E050UC; Perkin-Elmer) diluted 2:3 (v/v) with 2 mM cold lysine) to a final lysine concentration of 12 µM. At the desired time points, 100 ul of reaction was passed over a Dowex cation exchange resin column (N-methyl D-glucamine form) (217484; Millipore-Sigma) to remove any uninternalized ^14^C-lysine. The eluted proteoliposomes were resuspended with scintillation fluid (6013111; Perkin-Elmer) for liquid scintillation counting.

### *In vivo* amino acid starvation and microscopy

For *in vivo* experiments, Ypq1-GFP was expressed under its native promoter in yeast auxotrophic to multiple amino acids (YML1415). Cells were grown in rich media (YPD) in 26°C overnight with shaking. The next day, cell pellets were centrifuged at 13,500xg for 1 min, washed twice with MQ H_2_O, transferred to YNB media lacking the test amino acid, and incubated at 26°C for 6 hrs with shaking. At time intervals, cells were collected for 1 min at 13,500xg, washed with MQ H2O, and imaged immediately.

Microscopy was performed with a DeltaVision Elite system (GE Healthcare Life Sciences), equipped with an Olympus IX-71 inverted microscope, a scientific Complementary Metal-Oxide Semiconductor (sCMOS) camera, a 100×/1.4 Oil Super-Plan Apo-chromatic objective, and a DeltaVision Elite Standard Filter Set with the FITC filter (Excitation:475/28, Emission:525/48) for GFP. Image acquisition and deconvolution were performed in the program Softworx. ImageJ (NIH) was used for image processing.

### Western blot

For starvation experiments, starvation and Western blot were performed based on established protocols (Arines and Li, 2022; Yang et al., 2021). Briefly, cells expressing Ypq1-GFP under its native promoter were grown to mid-log (OD600: 0.5-0.8) in YPD media at 26°C with shaking. Cells were collected at 13,500xg for 5 mins, washed twice with MQ H_2_O, and incubated in YNB media lacking the test amino acid at 26°C with shaking. At time intervals, 7 OD cells were collected at 13,500xg for 5 mins and incubated on ice for at least 30 min in 10% TCA, followed by one wash of 0.1% TCA. Cell lysates were prepared by bead-beating for 5 min in 2x urea boiling buffer (50 mM Tris, pH 7.5, 10 mM EDTA, pH 8.0, 6 M urea, 200 mM DTT, 4% w/v SDS) and then in 2x urea sample buffer (150 mM Tris, pH 6.8, 6 M urea, 100 mM DTT, 4% w/v SDS, 10% v/v glycerol, 0.01% w/v bromophenol blue), incubating samples for 5 min at 42°C after every bead-beating step. Cell lysates were centrifuged at 14,000 g for 5 min, and the supernatant was collected. Samples were separated via electrophoresis on 11% polyacrylamide gels, then transferred to nitrocellulose membranes. Membranes were probed with the primary antibodies rabbit anti-GFP (1:2,500; TP401; Torrey Pines Biolabs), mouse anti-Pgk1 (1:5,000; 459250; Invitrogen), and secondary antibodies green goat anti-rabbit (1:10,000; 926–32211; LI-COR Biosciences) and red goat anti-mouse (1:10,000; 926–68020; LI-COR Biosciences).

Using LI-COR Image Studio, the fluorescence signal of full-length (FL) Ypq1-GFP and free GFP were measured. FL Ypq1-GFP (%) was quantified by dividing FL Ypq1-GFP by the total GFP signal (FL Ypq1-GFP + free GFP).

**Table 1.**
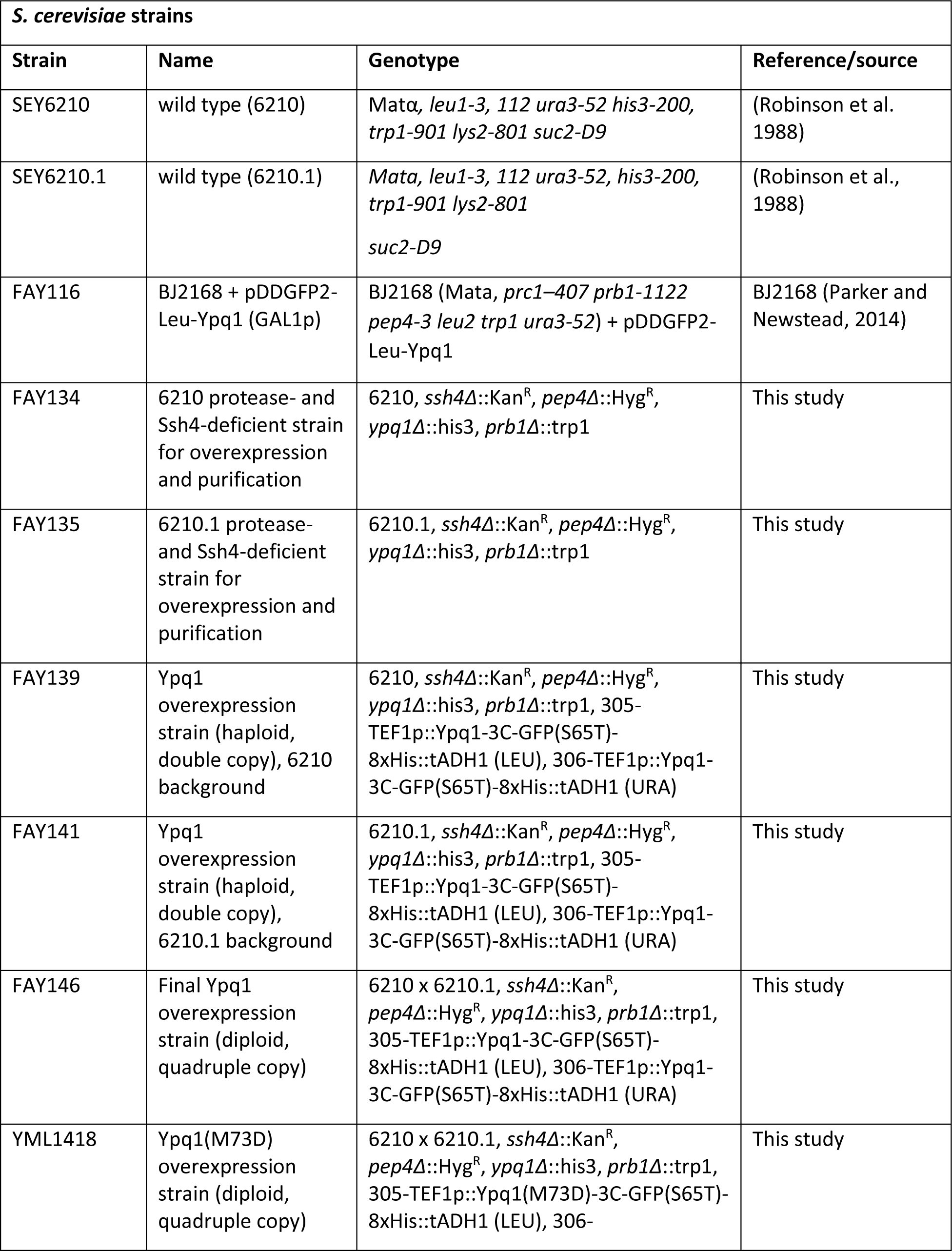
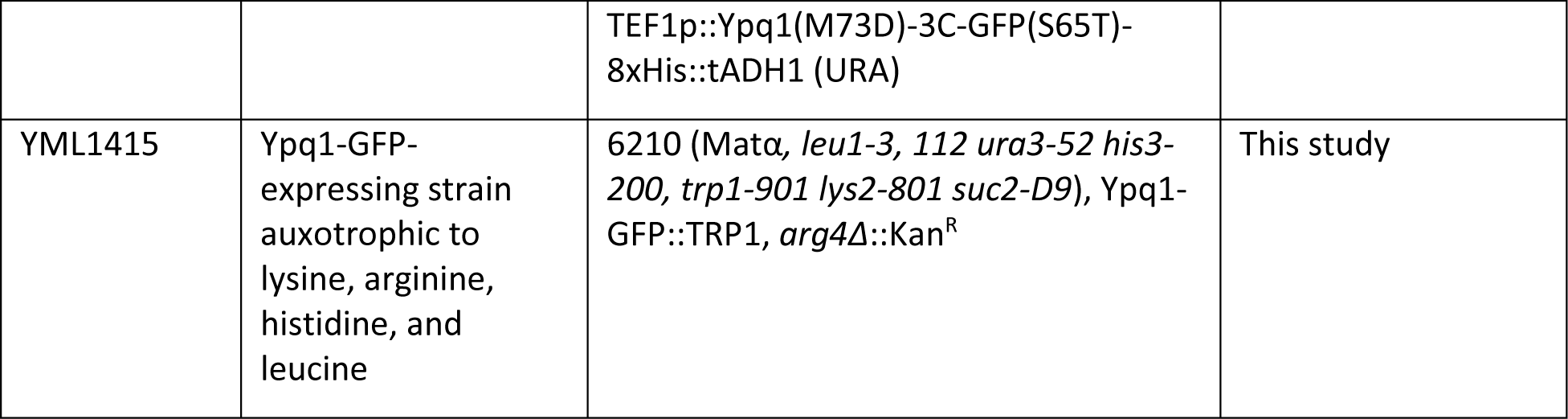
List of yeast strains used in this study.

**Table 2.** Yeast expression plasmids used in this study.

| ***S. cerevisiae* expression plasmids** |  |  |  |  |
| --- | --- | --- | --- | --- |
| **Code** | **Vector** | **Name** | **Description** | **Reference/source** |
| MLP857 | pRS426 (2 $\mu$, *URA3 LEU2 double selection*) | pDDGFP2-Leu2 | pRS426-GAL1p::TEV-yeGFP-8xHis | (Parker and Newstead, 2014) |
| PFA380 | pRS426 (2 $\mu$, *URA3 LEU2 double selection*) | GAL1p::Ypq1 overexpression plasmid (*URA3 LEU2*) | pRS426-GAL1p::Ypq1-TEV-yeGFP-8xHis with additional Leu selection | This study |
| PFA390 | pRS305 (*LEU2 integration*) | TEF1p::Ypq1-3C-GFP(S65T)-8xHis integration plasmid (*LEU2*) | pRS305-TEF1p::Ypq1-3C-GFP(S65T)-8xHis::tADH1 | This study |
| PFA391 | pRS306 (*URA3 integration*) | TEF1p::Ypq1-3C-GFP(S65T)-8xHis integration plasmid (*URA3*) | pRS306-TEF1p::Ypq1-3C-GFP(S65T)-8xHis::tADH1 | This study |
| PFA396 | pRS305 (*LEU2 integration*) | TEF1p::Ypq1 PQ mutant integration plasmid (*LEU2*) | pRS305-TEF1p::Ypq1 (P229S,Q230R)-3C-GFP(S65T)-8xHis::tADH1 | This study |
| PFA398 | pRS306 (*URA3 integration*) | TEF1p::Ypq1 PQ mutant integration plasmid (*URA3*) | pRS306-TEF1p::Ypq1 (P229S,Q230R)-3C-GFP(S65T)-8xHis::tADH1 | This study |
| PFA397 | pRS305 (*LEU2 integration*) | TEF1p::Ypq1 M73D mutant integration plasmid (*LEU2*) | pRS305-TEF1p::Ypq1(M73D)-3C-GFP(S65T)-8xHis::tADH1 | This study |
| PFA399 | pRS306<br>(*URA3*<br>*integration*) | TEF1p::Ypq1 M73D<br>integration plasmid<br>(URA3) | pRS306-TEF1p::Ypq1(M73D)-<br>3C-GFP(S65T)-8xHis::tADH1 | This study |

## Acknowledgments

We thank members of the Li laboratory for their helpful discussion and technical support. We thank B. McIlwain and T. Yeh from the Stockbridge lab for their biochemical expertise and technical assistance. We thank Dr. Liang Feng from Stanford School of Medicine for sharing the reagents. F. Arines was supported by the Barbour Scholarship from the University of Michigan Rackham Graduate School. This research is supported by National Institutes of Health grants (R01GM133873 to ML and R35GM128768 to RBS).

## Author contributions

Conceptualization, FA, RBS, and ML; Methodology, FA, OB, RBS, and ML; Investigation, FA, AW, FG, and ML; Writing & Editing, FA, RBS, and ML; Funding Acquisition, RBS and ML; Resources & Supervision, RBS and ML.

## Competing interests

The authors declare no competing interests.

**Fig. S1.**
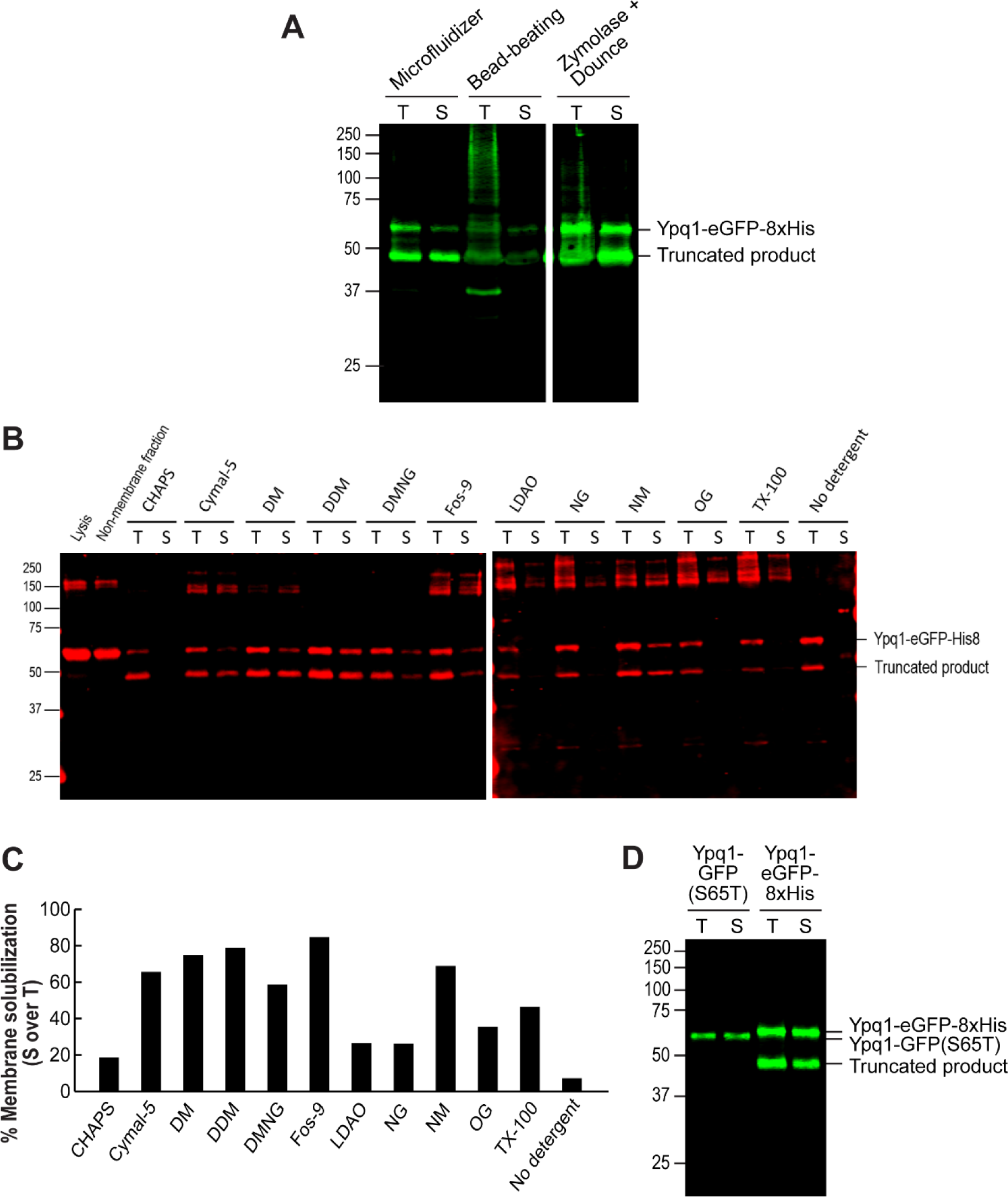
Optimization of purification conditions. Pellets of equivalent wet weights were solubilized in the corresponding detergents and centrifuged to pellet insoluble material. Total (T) vs. Solubilized (S) fractions were analyzed by Western blot. **A)** Comparison of different lysis methods on yield and solubility of Ypq1-GFP-8xHis. Pellets were solubilized in 0.5% DDM and centrifuged to pellet insoluble material. Fractions were probed using a GFP antibody. **B)** Eleven detergents at concentrations above CMC were used to solubilize Ypq1-GFP-8xHis. Fractions were probed using a His antibody. **C)** Quantification of bands in Solubilized fractions (Ypq1-eGFP-8xHis + Truncated product) over Total. Fos-9, DDM, and DM are the best solubilizers, while CHAPS, LDAO, and NG are the poorest solubilizers. We also took note of the appearance of high molecular-weight bands, which could correspond to aggregates. **D)** Comparison of GFP variants on fusion protein stability. Pellets were solubilized in 0.5% DDM and centrifuged to pellet insoluble material. Switching to GFP(S65T) eliminated the formation of truncation products.

